# ChAdOx1 COVID vaccines express RBD open prefusion SARS-CoV-2 spikes on the cell surface

**DOI:** 10.1101/2023.05.22.541685

**Authors:** Tao Ni, Luiza Mendonça, Yanan Zhu, Andrew Howe, Julika Radecke, Pranav M. Shah, Yuewen Sheng, Anna-Sophia Krebs, Helen M. E. Duyvesteyn, Elizabeth Allen, Teresa Lambe, Cameron Bisset, Alexandra Spencer, Susan Morris, David I. Stuart, Sarah Gilbert, Peijun Zhang

**Author notes:** These authors contributed equally to this study (Tao Ni, Luiza Mendonça, Yanan Zhu). Department of Biochemistry, Molecular Biology and Biophysics, University of Minnesota, Minneapolis, 55455, USA. School of Biomedical Sciences, LKS Faculty of Medicine, The University of Hong Kong, Pokfulam, Hong Kong SAR, China.

## Abstract

Vaccines against SARS-CoV-2 have been proven to be an effective means of decreasing COVID-19 mortality, hospitalization rates, and transmission. One of the vaccines deployed worldwide is ChAdOx1 nCoV-19, which uses an adenovirus vector to drive the expression of the original SARS-CoV-2 spike on the surface of transduced cells. Using cryo-electron tomography and subtomogram averaging, we determined the native structures of the vaccine product expressed on cell surfaces *in situ*. We show that ChAdOx1-vectored vaccines expressing the Beta SARS-CoV-2 variant produce abundant native prefusion spikes predominantly in one-RBD-up conformation. Furthermore, the ChAdOx1 vectored HexaPro stabilized spike yields higher cell surface expression, enhanced RBD exposure, and reduced shedding of S1 compared to the wild-type. We demonstrate *in situ* structure determination as a powerful means for studying antigen design options in future vaccine development against emerging novel SARS-CoV-2 variants and broadly against other infectious viruses.

## Introduction

The current COVID-19 pandemic provoked the fastest vaccine development effort in the history of mankind: approximately a year from disease report to vaccine distribution. Previously the record belonged to the mumps vaccine, which took about four years from initiation of development to deployment. The majority of WHO-recognized SARS-CoV-2 vaccines are based on the spike (S) glycoprotein. S decorates the exterior of viral particles and is responsible for both attachment of viruses to the target cells through its interaction with the ACE2 receptor and the fusion of the viral membrane with the host cell membrane to deliver the viral content to the target cell^1-3^. S is a homotrimer that undergoes massive structural rearrangements during the viral entry and fusion steps. The prefusion conformation, composed of S1 and S2 subunits, is metastable, active, and responsible for binding to the ACE2 receptor. The postfusion conformation is formed after shedding S1 and is more stable but inactive. Immune responses against the prefusion conformation are more likely to be protective, as they may impede viral entry and subsequent infection. The natural propensity of the spike to convert to the postfusion conformation may therefore present a challenge in vaccine development^4^. To address this issue, a structure-based design approach identified six proline substitutions (F817P, A892P, A899P, A942P, K986P, and V987P) in S2 (HexaPro) that increase stability and expression yield of the SARS-CoV-2 prefusion spike^5^.

One of the COVID-19 vaccines granted Emergency Use Licensure by the World Health Organisation is AstraZeneca’s ChAdOx1 nCoV-19/AZD1222, which is based on a replication-incompetent adenoviral vector^6^. This viral vector enters cells and delivers the gene for the SARS-CoV-2 spike, leading to its transient expression. AZD1222 drives expression of the spike from the Wuhan-Hu-1 strain (GenBank accession no. MN908947), an early SARS-CoV-2 isolate^7^. The spike derived from this vaccine adopts a similar glycosylation pattern and overall structure as the one found in native SARS-CoV-2 viruses^8^. Vaccination with AZD1222 triggers cellular and humoral immune responses that greatly reduce COVID-19 infection, deaths, and hospitalizations^6,9^. During the pandemic, a number of spike variants were expressed from ChAdOx1, including the beta variant both with (described here as 19E6) and without (described here as 19E, elsewhere as AZD2816) Hexapro stabilisation^10,11^.

## Results

We investigated the second-generation ChAdOx1 constructs against the Beta/South Africa SARS-CoV-2 variant, specifically the ChAdOx1 19E and 19E6. U2OS cells were transduced by each of the ChAdOx1 constructs. The expression levels of SARS-CoV-2 spike proteins were analyzed by flow cytometry and staining at three post-transfection time points (24h, 48h, and 72h) (Supplementary Fig. 1). ACE2-Fc was used to detect the functional spike which has at least one RBD up conformation while a monoclonal human antibody Ab222^12^ was used to probe the prefusion state of the spike. In both constructs, the expression level of spikes at 24h post-infection is high and does not further increase after longer time expression. 19E6 construct showed both a higher percentage of spike-producing cells (Fig. 1A-B) and a much higher level of surface expression (Fig. 1C-D) of the spike in the prefusion functional state than the 19E construct. The higher level of the surface spike in 19E6 could be attributed to the increased stability of the spike. Indeed, the level of S1 shedding in the extracellular media in the 19E6 is considerably lower than 19E, especially at 72h post-infection (Fig. 1E). Transfection of ChAdOx1 does not affect the viability of cells (Fig. 1F).

**Figure 1.**
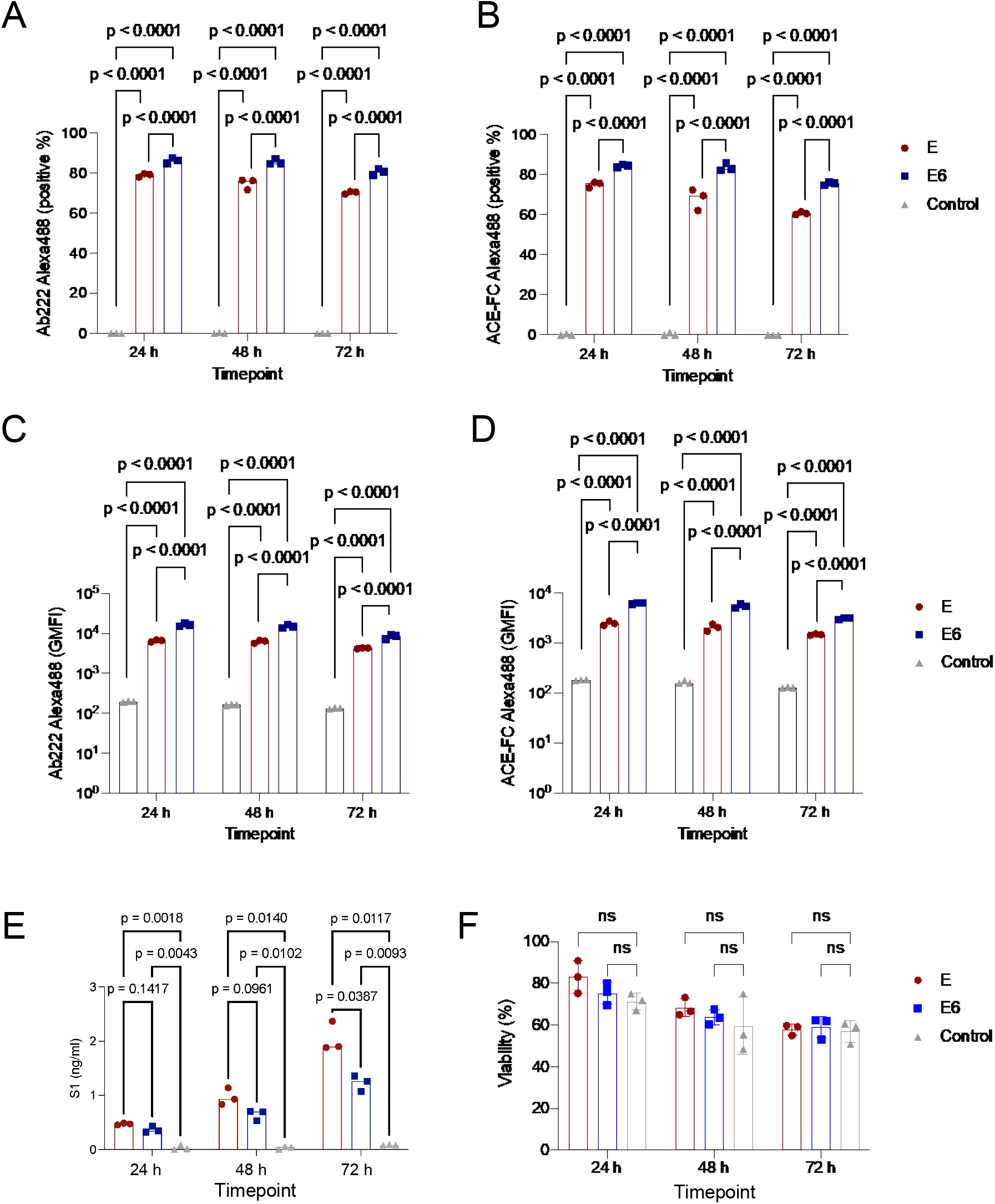
Cytometry analysis of SARS-CoV-2 spike expression. (A and B) Percentage of cells expressing SARS-CoV-2 spike after ChAdOx1 19E and 19E6 transduction as assessed by staining with Ab222, a S1-targetting monoclonal antibody (A) and an ACE2-Fc chimera (B). (C and D) Cell surface expression levels of SARS-CoV-2 spike after ChAdOx 19E and 19E6 transduction as assessed by staining with Ab222 (C) and a ACE2-Fc chimera (D). (E) S1 concentration in microvesicle-depleted cell-culture supernatant determined by quantitative S1 ELISA. (F) Viability of cells transfected with ChAdOx1 19E and 19E6 at 24h, 48h and 72h post infection. p values from two-way ANOVA with Tukey multiple comparisons test. Experiment done two times with biological triplicates.

To understand what conformation(s) the vaccine spikes adopt on transduced U2OS cell surfaces, we collected cryoET tilt-series to image the cell periphery (Supplementary Fig. 2, Supplementary Table 1). As shown in representative tomograms (Fig. 2A and Movie 1-2), a remarkably high amount of spike protein is seen on the cell surface, as seen by the club-head shapes in the side view and triangular shapes in the top view (Fig. 2A). Spike-covered exosomes are also commonly identified (Supplementary Fig. 2B). These spikes are packed on the membrane surfaces with the same height, resembling hedgerows, suggesting they are mostly prefusion spikes. We performed cryoET subtomogram averaging (STA) of these *in situ* spikes and obtained the maps with C3 symmetry at 9.0 Å and 9.6 Å resolution for the 19E6 and 19E construct, respectively (Fig. 2B and Supplementary Fig. 3A). Local resolution maps suggest a stable central core and much more dynamic RBD domains (Fig. 2B). The central helix bundles are resolved clearly (Fig. 2C), as well as a number of N-linked glycans (Fig. 2D). The majority of spikes on the native cell membrane are mono-dispersed with a median distance of ∼15 nm as shown in the nearest neighbor distance distribution, distinct from the dimer configuration observed in detergent-solubilized and purified recombinant spikes^13^ (Fig. 2E-F).

**Figure 2.**
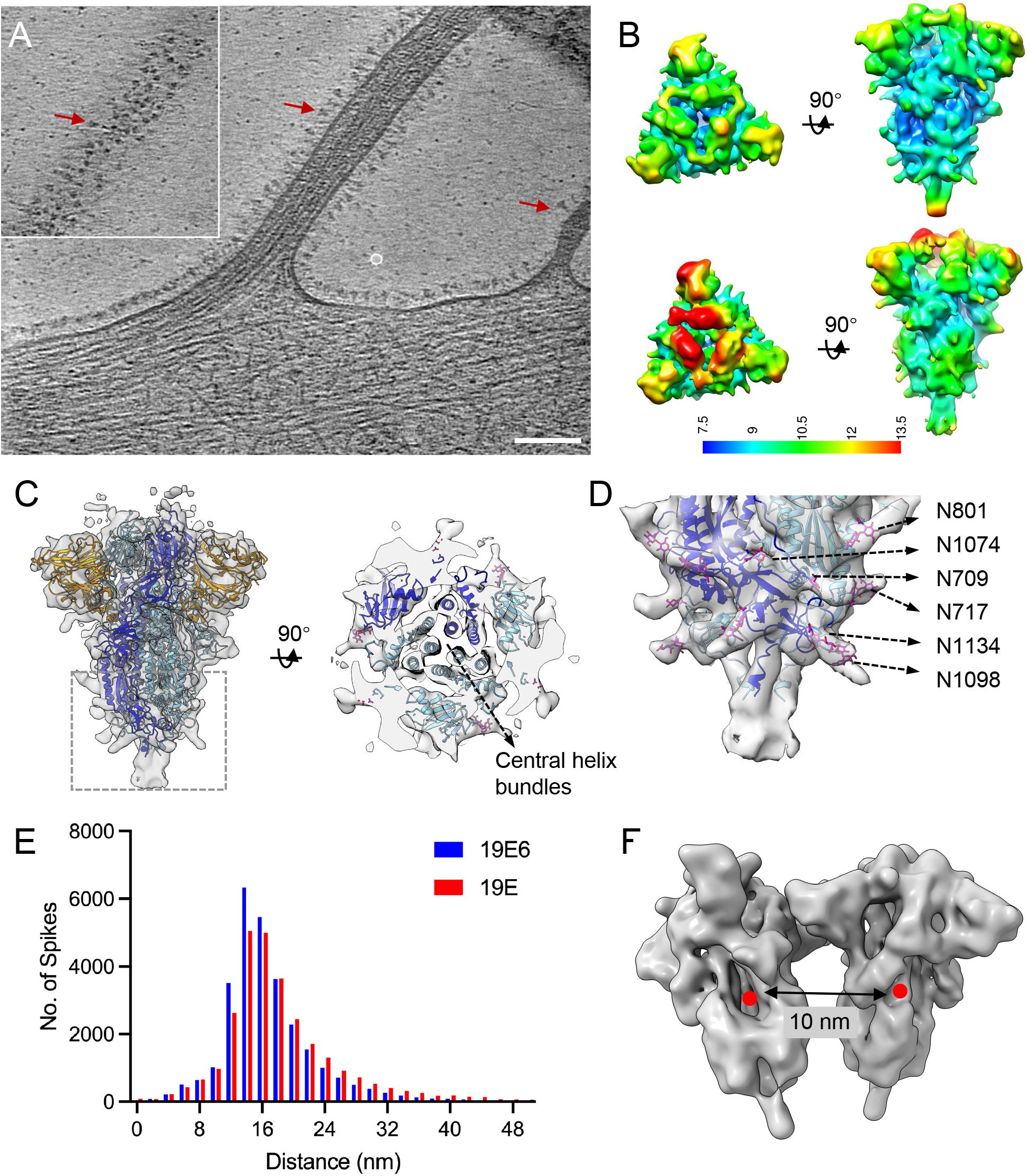
Subtomogram averaging of ChAdOx1 spikes *in situ*. (A) A tomogram slice of 19E6 infected cell showing spikes decorating the cell membrane. Red arrows point to the representative spikes. Inset box shows the spikes in the top slide of the filopodia. Scale bar = 100 nm. (B) Orthogonal views of subtomogram averaging of ChAdOx1 spike 19E6 (top) and 19E (bottom) with C3 symmetry applied. (C) Orthogonal views ChAdOx1 spike 19E6 density map overlay with a structure model of spike (PDB 6ZGG). (D) A close-up view of the boxed region in C. The density extrusions indicate good fit with the N-linked glycosylation. Several glycans are highlighted. (E) Pairwise distance distribution of spikes for ChAdOx1 19E and 19E6 samples. (F) Model of dimerized spikes (EMD-22354) has a distance of 10 nm between spikes. The density map EMD-22354 was gaussian filtered for presentation.

The poorer local resolution in the RBD domain indicates that it may be conformationally flexible (Fig. 2B). We, therefore, carried out subtomogram classification without symmetry in emClarity^14^ using a PCA-based reference-free classification method (Supplementary Fig. 4). This analysis revealed that the majority of spikes are in the one-RBD-up conformation and smaller fractions of spikes are in all-RBD-down or two-RBD-down with the third RBD missing/flexible conformations (Fig. 3A). The same three distinct classes of spikes at a similar ratio were reproduced using a reference-based classification (Supplementary Fig. 4). Intriguingly, the distribution of the three classes for the 19E6 spikes is markedly different from the 19E spikes (Fig. 3B). More than 85% of 19E6 spikes adopt one-RDB-up conformation as opposed to 58% of 19E spikes. Previous structural analyses of spikes from inactivated virions and from recombinant soluble proteins showed multiple spike conformations with the majority of the spikes in one-RBD-up conformation^15^, supporting the conformational authenticity of the antigen generated by the ChAdOx1 vaccine, both with and without HexaPro stabilization. Further refinement of the one-RBD-up spike class resulted in a structure at 9.6 Å for the 19E6 spike which closely resembles the purified spike from *in vitro* studies (Fig. 3C-D).

**Figure 3.**
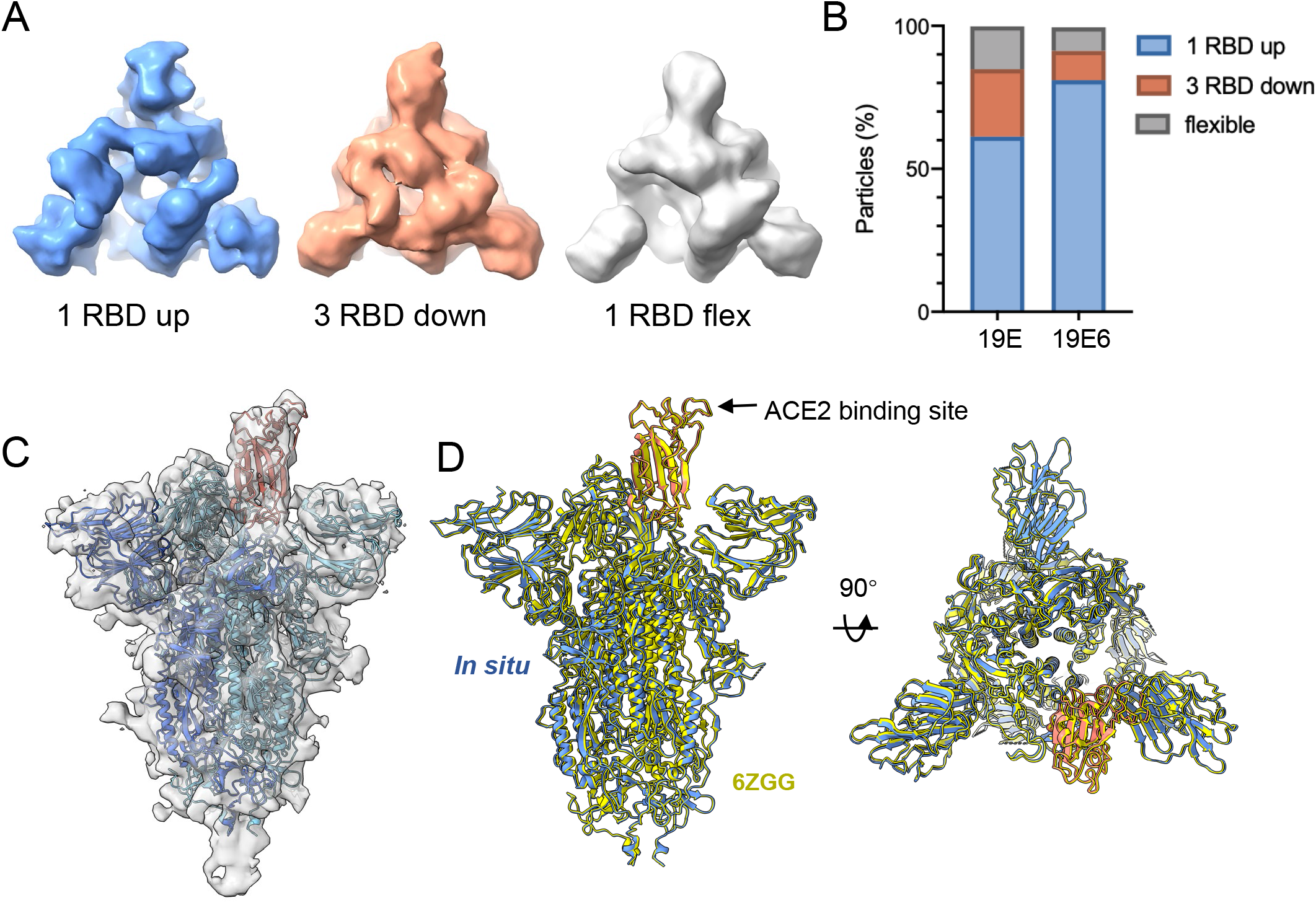
Classification of spike conformations. (A) Three classes were identified in both ChAdOx1 19E and ChAdOx1 19E6 spikes: one RBD up (blue), three RBD down (brown) and two RBD down with one RBD flexible (grey). (B) Distribution of the three spike conformations in both ChAdOx1 19E and ChAdOx1 19E6 vaccines. (C) STA map of ChAdOx1 19E6 spike in one RBD up conformation at 9.6 Å resolution. (D) Comparison between ChAdOx1 19E6 spike (one RBD up) in situ with cryo-EM SPA study (PDB 6ZGG) with a RMSD of 0.875 Å. The ACE2 binding site is on the top of RBD domain.

## Discussion

We determined native structures of the vaccine products in the cellular context by cryoET, allowing vaccine characterization and validation. We compared two second-generation ChAdOx1 constructs expressing the wild-type and HexaPro spikes of the Beta SARS-CoV-2 variant and revealed that the HexaPro ChAdOx1 spikes are more stable, with enhanced cell surface expression, improved RBD exposure, and reduced S1 shedding. While the original *in vitro* study of the recombinant HexaPro mutant suggested an approximately equal distribution of spikes in one-RBD-up, two-RBD-up, and one-RBD-flexible prefusion state^5^, our *in situ* structure from the cell surfaces, showed that the spikes are mostly in the prefusion state with about 85% of spikes in one-RBD-up conformation. HexaPro stabilized SARS-CoV-2 spikes delivered by a Newcastle Disease Virus vector are currently being used in a clinical phase 1 trial. Interim analyses show an acceptable safety profile and potent immunogenicity^16^. In mice and hamsters, HexaPro spike delivered by a Vesicular Stomatitis Virus vectored vaccine is more potent for inducing antibodies that neutralize viral variants of concern than the 2-Proline mutant and the wild-type spike^17^. However, in pre-clinical studies, vaccination of BALB/c mice with the ChAdOx1 vectors expressing Beta variant HexaPro-stabilized spikes were shown to induce equivalent IgG responses to non-stabilized spikes and induced slightly lower interferon-gamma production in splenocytes when compared to non-stabilized spikes, suggesting a moderately lower cell-mediated immune response^17^. The complexity of neutralizing antibody induction and immune cell responses by various virus-vectored vaccines in different hosts warrants further in-depth investigations but may relate to the different expressions of the vaccine antigen from VSV and ChAdOx1. VSV-vectored vaccines carry the vaccine glycoprotein on the surface of VSV particles, and the spike protein is therefore present during vaccine manufacture and storage, perhaps allowing time for the antigen to alter conformation from the prefusion to postfusion form. In contrast, high levels of vaccine antigen are produced in cells transduced with ChadOx1 only after vaccination, with a significant amount of the antigen remaining in the prefusion form. However, the addition of prolines may adversely affect the processing of the vaccine antigen by the proteasome, thereby reducing T-cell responses to the antigen.

The continuous emergence of SARS-CoV-2 variants will most likely require regular booster vaccinations to keep up the arms race against the virus. Optimizing antigens in future-generation vaccines will be crucial in the fight against SARS-CoV-2 and to prepare for potential future coronavirus outbreaks. Structural characterization and validation *in situ* are paramount for these vaccine candidates, especially when the antigenicity of a vaccine antigen is not predictive of the protective immunity elicited by it, as the FI-RSV trial shows^18^. Our *in situ* structural study involves minimal experimental manipulation of spikes, which reflects the closest recapitulation of antigen status presented in their native condition. These structural and conformational characterizations of spikes *in situ* provide a basis for the development of next-generation immunogens, meeting the different requirements of alternative vaccine technologies.

## Data availability

All data needed to evaluate the conclusions in the paper are present in the paper and/or the Supplementary information. The motion-corrected tilt-series have been deposited to EMPIAR with the accession codes as follows: ChAdOx1 19E6, EMPIAR-11456, and ChAdOx1 19E, EMPIAR-11457. The cryo-ET subtomogram averaging density maps have been deposited in the EMDB with the accession codes as follows: ChAdOx1 19E6 spike C3 symmetrized (EMD-16403); ChAdOx1 19E6 spike in C1 symmetry (EMD-16404, all particles); ChAdOx1 19E6 spike in C1 symmetry (EMD-16697, one-RBD-up only); ChAdOx1 19E spike C3 symmetrized (EMD-16405); ChAdOx1 19E spike in C1 symmetry (EMD-16406). The MATLAB scripts have been uploaded to Github (https://github.com/Tao-Ni/emClarity_scripts.git).

## Acknowledgments

We acknowledge Diamond for access and support of the cryo-EM facilities at the U.K. national eBIC (proposal BI26987), funded by the Wellcome Trust, MRC, and BBSRC. We acknowledge The Oxford Particle Imaging Centre (OPIC) for access to the Glacios which was founded by a Wellcome JIF award (060208/Z/00/Z) and is supported by a Wellcome equipment grant (093305/Z/10/Z). This research was supported by the UK Wellcome Trust Investigator Award (206422/Z/17/Z) and the ERC AdG grant (101021133), and by the Wellcome Trust Core Award Grant Number 203141/Z/16/Z with additional support from the NIHR Oxford BRC, MRC (MR/N00065X/1) and the Chinese Academy of Medical Sciences (CAMS) Innovation Fund for Medical Science (CIFMS), China (2018-I2M-2-002). Additional support was received from AstraZeneca.

## Contributions

P.Z. conceived the project. P.Z., D.S., and S.G. designed experiments and data analyses. L.M., E.A., A.S., T.L., C.B., and S.M. carried out the Flow cytometry and ELIZA experiments and analysis. L.M. and A-S.K. prepared the transduced cells and made cryo-EM grids. L.M. and H.D. screened grids. A.H., J.R., T.N., and Y.Z. collected cryo-ET datasets. T.N. performed cryo-ET subtomogram averaging with the assistance of Y.Z., Y.S., A-S.K., and P.S.. T.N. performed cryo-ET subtomogram classification and analyzed the structures. T.N., L.M., and P.Z. wrote the manuscript with support from all authors.

## Online methods

### Production of ChAdOx1 19E and 19E6

The construction of ChAdOx1 nCov-19 expressing the glycoprotein (S) gene from from strain B.1.351 of SARS CoV-2 first identified in South Africa, as well as the derived vector containing 6 proline substitutions (F817P, A892P, A899P, A942P, K986P, and V987P) has been described previously^11^. Briefly, the sequence was codon-optimized for expression in human cell lines and synthesized with the tissue plasminogen activator (tPA) leader sequence at the 5′ end by GeneArt Gene Synthesis (Thermo Fisher Scientific). The S genes were inserted into the Gateway® recombination cassette of the shuttle plasmid containing a human cytomegalovirus major immediate early promoter (IE CMV), which includes intron A and two tetracycline operator 2 sites, and the bovine growth hormone polyadenylation signal. BACs containing the ChAdOx11 SARS CoV-2 S were prepared by Gateway® recombination between the ChAdOx11 destination DNA BAC vector^19^ and the shuttle plasmids containing the SARS CoV-2 gene expression cassettes using standard protocols resulting in the insertion of the SARS-CoV-2 expression cassette at the E1 locus. The ChAdOx1 SARS CoV-2 S adenovirus genome was excised from the BAC using unique PmeI sites flanking the adenovirus genome sequence. ChAdOx1 SARS CoV-2 S viral vectors were rescued in T-REx TM cells (Invitrogen, Cat. R71007). The resultant viruses, ChAdOx1 nCov-19E and ChAdOx1 nCoV-19E6, were purified by CsCl gradient ultracentrifugation as described previously^20^. The infectious titers were determined on T-REx TM cells using an anti-hexon immunostaining assay based on the QuickTiter™ Adenovirus Titer Immunoassay kit (Cell Biolabs Inc).

### Electron Microscopy Grid preparation

Gold G300F1 grids with Quantifoil R2/2 carbon were glow discharged (Harrick Plasma) on the highest setting for 45 seconds, transferred carbon-side up to a 6-well plate, and treated with 20 mg/ml of Bovine Plasma Fibronectin (Sigma) in PBS for 30 minutes and washed three times with PBS. Grids were then UV-treated using a short-wave mercury lamp (Jena Analitik) for 1 h. U2OS cells were seeded in 6 well plates on top of the EM grids. 3 to 6×10^4^ cells were used in each well. Plates were incubated at 37°C overnight before ChAdOx1 SA 19E and ChAdOx1 SA 19E6 transduction. The ChAdOx1 vector was diluted in serum-free media and an MOI of 10 was used for all transductions. Cell media was replaced by serum-free media and the vector dilutions were added drop-wise to each well. Cells were incubated at 37°C for 4 h before the addition of serum-containing media to a final concentration of 10% FBS. Non-transduced cell controls were performed for each time point with the same conditions. Plates were incubated at 37°C until plunge-freezing. Grids were picked up from the wells and gently washed three times with PBS. A few microlitres of concentrated 10 nm Gold fiducials (EMS) were applied to the gold side of grids, which were then blotted on the gold side and immediately plunge-frozen in liquid ethane using a Leica GP2 plunge freezer. Grids were stored in liquid nitrogen until cryo-ET imaging. The remaining cells on the 6-well plates were used for FACS and viability analyses.

### Cell viability analysis

Cells were detached from the wells using a cell scraper and resuspended in Phosphate Buffered Saline (PBS). 20 ml of cell suspension was mixed with 20 ml of 0.4% Trypan Blue solution (Sigma) and manually counted using a hemocytometer. Each sample was counted in technical duplicate. Three biological replicates were performed for each condition.

### Flow Cytometry

ChAdOx1-transduced and non-transduced control cells at 24, 48, and 72 h post-transduction were detached from the wells using a cell scraper and resuspended in PBS. Cells were centrifuged at 1500 ×g for 5 minutes and washed with 1% BSA in PBS before being incubated with 1 mg/ml human monoclonal Ab222 or 2 mg/ml recombinant human ACE2-Fc for 2 h at room temperature. Cells were washed three times using 1% BSA in PBS and fixed using 1% paraformaldehyde in PBS for 30 minutes at room temperature. Cells were washed three times with 1% BSA in PBS and then incubated for 1 h at room temperature with Goat Anti Human AlexaFluor 488 secondary antibody (Life Technologies). Cells were washed twice and then analyzed by flow cytometry using a Fortessa X20 FACS analyzer. Cells were considered positive for spike expression if they had a fluorescence intensity above a threshold value determined by the maximum intensity of the non-infected control cells. Two experiments were done independently, with three biological replicates performed for each condition. Data were analyzed using FlowJo v9 (TreeStar).

### S1 ELISA

The supernatant was harvested from ChAdOx1-transduced and non-transduced control cells at 24, 48, and 72 h time points. They were spun at 1000 ×g for 5 min and filtered through a 0.45 μm syringe filter. S1 concentration was assessed using a SARS-CoV-2 Spike Protein S1 RBD ELISA Kit (Catalogue No. E-EL-E605-ELA, eLabScience) according to manufacturer instructions. Three biological replicates were performed for each condition.

### CryoET imaging

Tilt-series were acquired using ThermoFisher Titan Krios microscopes operated at 300 kV, equipped with a K3 camera and Quantum energy filter in zero-loss mode. The energy slit width was set to 20 eV. The tilt series were collected with serialEM using a dose-symmetric scheme, starting from 0° with a 3° tilt increment by a group of 3, with an angular range from -60° to 60°. The accumulated exposure for each tilt series was ∼120 e-/Å^2^, with a defocus range between -2 and -5 μm and a pixel size of 2.2 Å. Ten raw frames without gain normalization at each tilt were saved in tif format and data collection details are listed in Supplementary Table 1.

### Subtomogram averaging

An in-house on-the-fly script toolbox.py was used to generate tilt-series and tomograms for visualization, from raw micrographs (https://github.com/ffyr2w/cet_toolbox). Briefly, the raw micrographs were gain normalized and motion-corrected using MotionCor2, removing the first frame in the raw tilt. The images were stacked together using the *newstack* command in IMOD and aligned with *batchruntomo* using gold fiducial beads. The aligned tilt-series were manually checked in Etomo and only high-quality tilt-series were included for further subtomogram averaging: (1) > 5 gold fiducial beads for tilt-series alignment; (2) visible spikes on the membrane from the tomogram reconstructions. Tilt series with no identifiable spikes were excluded for further processing.

Subtomogram averaging of the two datasets was performed separately using the same pipeline and parameters in emClarity^14^. First CTF and defocus gradient were estimated. The spikes on the membrane were then identified through template matching in emClarity (v1.5.0.2) with none-CTF convoluted tomograms (4x down-sampled, pixel size of 8.8Å), using a prefusion spike (EMD-21452, closed state) filtered to 20 Å resolution as a template. The position and orientation of each spike were manually checked in Chimera with the Place Object plugin (v2.1.0)^21^. An in-house script (em2emClarity.m) was developed to convert emClarity database into pyTOM format for the Place Object plugin in Chimera. Particles in the wrong orientation or position with respect to the membrane were removed manually. The remaining particles were imported to emClarity for further processing (em2emClarity.m). The two scripts for converting files between emClarity database and pyTOM formats have been deposited to Github (https://github.com/Tao-Ni/emClarity_scripts.git).

The subtomograms were iteratively aligned and averaged using 3×, 2× and 1× down-sampled tomograms, initially with C3 symmetry applied. The rotational search ranges were iteratively reduced for each down-sampled tomogram until the resolution of averages did not further improve. The same dataset was also processed with C1 symmetry using 2× down-sampled tomograms, allowing a full-range in-plane rotational search after relaxation from the previous C3 symmetry alignment. Further local alignment was performed using 1x binned tomograms. The final half maps were reconstructed with cisTEM within the emClarity package with either C3 or C1 symmetry using the total exposure. One round of local translational search on the extracted projection images was performed to further improve the resolution of final reconstructions. The final combined maps with FSC weighting and B-factor sharpening were generated in RELION (v4.0)^22^. Local resolution estimation of density maps was conducted in RELION using relion_postprocess with a soft molecular mask.

### Subtomogram classification

Subtomogram classification was performed using emClarity^14,23^. Two different classification methods were applied: the standard PCA-based 3D classification and reference-based 3D classification. For the PCA-based 3D classification, the subtomograms were firstly aligned with C3 symmetry, then relaxed to C1 symmetry allowing ±120° rotation, with a 120° step size using 2x binned subtomograms (Figure S4A-B). Several iterations were allowed for the in-plane rotational alignment until convergence. The resulting structure was further aligned with local search before 3D classification (Figure S4C). For classification, a spherical mask enclosing the bulky domain of NTD and RBD trimer was used. 16 class averages were generated and further merged into three classes: three-RBDs-down, one-RBD-up, and one-RBD-flexible (Figure S4D-E). For reference-based classification, three references (all RBDs down, one RBD up, and one RBD disordered) were generated using *molmap* command in Chimera from structure models (PDB codes: 6ZGI, 6ZGG and 6ZGH), filtered at 15 Å resolution. The resulting maps were used as references in emClarity to calculate the cross-correlation coefficients (CCCs) against each subtomogram, allowing a minimal rotational and translational search. Each subtomogram was then assigned to one of the three classes based on its highest CCC. Details of the classification procedure are illustrated in Figure S4F.

### Model fitting

The 3-fold symmetrized density maps for both E and E6 reveal the feature of 3RBD down conformation. An initial rigid-body fitting using a spike structure from cryoEM SPA with all 3 RBDs down conformation (PDB 6ZGI) was attempted but did not fit well, due to the local rotation of the NTD. In contrast, rigid-body fitting of spike structure with one RBD up conformation (PDB 6ZGG) is better. Therefore, a single chain in RBD down conformation (PDB 6ZGG, chain A) was extracted and a rigid body fit into the density map; the other two chains were generated by rigid-body docking of the Chain A structure into density. For the C1 density map, the whole spike trimer with one RBD up conformation (PDB 6ZGG) was rigid-body fitted into the density map, and each chain was further rigid-body refined. Figures were prepared in ChimeraX^24^ and Chimera^25^.

### Quantification and Statistical Analyses

Statistical analyses were performed in GraphPad Prism 9. Two-way ANOVA with Tukey multiple comparisons test was performed to compare 19E, 19E6, and control FACS and cell viability results. The distances of the nearest spike neighbor were calculated using the coordinates of refined spikes in their corresponding tomograms. The center of the resulting subtomogram averaging map was used as the center of the spike. The distance between two spikes in the dimer of the spike trimers map (EMD-22354) was calculated using the same spike center. The histogram of distance distribution was plotted in GraphPad Prism9.

